# Unveiling the PET plastics degradation potential of the thermostable EstS1 esterase through integrated biochemical, structural, and morphological analyses

**DOI:** 10.1101/2025.10.14.682280

**Authors:** Shalja Verma, Divya Aggarwal, Meet Ashar, Anand Kumar Pandey, Anup Sutradhar, Sakshi Pandey, Debabrata Sircar, Jitin Singla, Pravindra Kumar

## Abstract

Enzymatic polyethylene terephthalate (PET) plastic degradation is a promising approach to combat the exploding plastic pollution. EstS1, a pH-tolerant, thermostable esterase, has been previously recognized for its degradation potential against phthalate diester plasticizers. The present study elucidates the exceptional potential of this enzyme to degrade crystalline PET plastic and its primary intermediate, bis(2-hydroxyethyl)terephthalate (BHET), into terephthalate. Kinetic analyses revealed that EstS1 degrades 75% of BHET in 1h, liberating mono(2-hydroxyethyl) terephthalate (MHET) and terephthalate as end products. The co-crystal structure of wild-type EstS1 with BHET exhibited the electron density of BHET, MHET, and ethylene glycol, including MHET bound at the active site, in a canonical tetrahedral intermediate conformation. The complex structure of BHET with the Ser154Ala mutant of EstS1 further accommodated two BHET molecules, one interacting directly with the catalytic triad and the oxyanion hole. MD simulation analysis revealed highly stable interactions of BHET at the active site of EstS1. Moreover, SEM imaging displayed significant degradation of the crystalline PET plastic film by EstS1 esterase over a period of 15 days, both under controlled and soil-based fluctuating environmental conditions, highlighting its versatility to varying environmental conditions. XPS analysis discovered the increase in -C-O-, -C-N-, -N-H-, and -N=O- bonds at the surface of EstS1-treated PET film, indicating effective degradation. Consequently, this comprehensive kinetic, structural, and morphology-based analysis of the PET-degrading potential of EstS1 esterase encourages further enzyme engineering studies to exploit the dual potential of EstS1 esterase to degrade both plastic and plasticizers.

## 1. Introduction

Polyethylene terephthalate (PET) plastic production reached 28 million metric tons in 2024 and is expected to grow to 35 million metric tons by 2030. The consistent increase in the production of PET plastic and its accumulation in different ecosystems is threatening the survival and adaptability of ecosystems (Jovanović *et al*., 2025). The micro-sized particles of PET plastics have also been identified in human tissues, including the brain, lungs, kidneys, heart, and placenta, from where they can pass to the fetus after crossing the placental barrier. The adverse effects of PET plastic accumulation inside the human body include disruption of the endocrine system, neuronal dysfunctions, heart strokes, renal toxicity, and developmental problems in the fetus (Enyoh *et al*., 2023; Sheriff *et al*., 2025). Although extensive efforts have been directed to reduce the generation of PET plastic waste and enhance recycling practices to alleviate the increasing plastic pollution, they are unable to meet the tons of plastic accumulated around the globe (Massoud & Dsilva, 2024). Further, the recalcitrant nature of crystalline PET plastic and the toxic residues released during chemical and physical processes limit the degradation of plastics. Biodegradation of PET plastic has been studied extensively using diverse microbial strains or their respective enzymes (Maurya *et al*., 2020).

Serine hydrolases, which include cutinases, lipases, and esterases, are known to mediate the depolymerization of PET into bis(2-hydroxyethyl)terephthalate (BHET), which is subsequently degraded into mono(2-hydroxyethyl) terephthalate (MHET). MHET is further converted into terephthalic acid (TPA) and ethylene glycol, either by the same or another serine hydrolase (Chen *et al*., 2018) (Figure 1). These hydrolases contain a catalytic triad of the Ser-His-Asp/Glu that cleaves the ester bond found in PET or its degradation products using a ping-pong mechanism combined with tetrahedral covalent enzyme intermediates (Lai *et al*., 2023). Thermostability is a highly desired characteristic for PET depolymerizing hydrolases due to the high glass transition temperature of PET, lying in the range of 70 to 80 °C (Zhang, 2022). Several thermostable PET-degrading hydrolases have been identified, including *Ca*PETase, PHL7, *Tf*Cut2, *HiC* cutinase, Cut190, Est119, and LCC (Hong *et al*., 2023; Richter *et al*., 2023; Mrigwani *et al*., 2022; Ronkvist *et al*., 2009; Numoto *et al*., 2020; Kitadokoro *et al*., 2012; Sulaiman *et al*., 2011). Recent studies have further used directed evolution or AI/ML-based approaches to engineer or design thermostable or high-efficiency hydrolases for effective PET degradation, such as HotPETase, TurboPETase, DepoPETase, and PES-H1^L92F/Q94Y^ (Bell *et al*., 2022; Cui *et al*., 2024; Shi *et al*., 2023; Pfaff *et al*., 2022).

**Figure 1:**
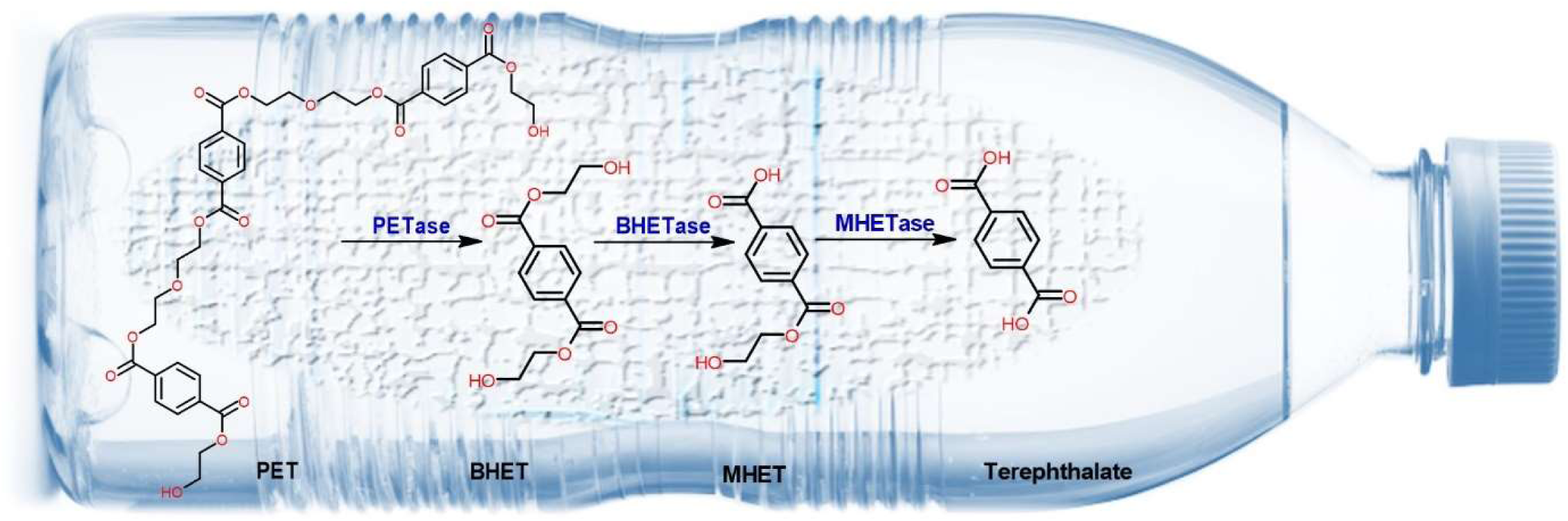
Enzymatic degradation of Polyethylene terephthalate (PET) into BHET, MHET and Terephthalate.

Although all these hydrolases reveal effective PET degradation, their efficiency has been mainly demonstrated to depolymerize amorphous PET film or powder of amorphous PET, but rarely crystalline PET (Arnal *et al*., 2023). As the PET plastic waste consists of high-crystallinity PET plastic and its products, an efficient thermostable PET-degrading hydrolase capable of degrading crystalline PET sheets is of immense importance (Thomsen *et al*., 2023). Moreover, plastics, after being discarded, end up in the soil and water bodies of different climatic conditions, which contain different compositions of ionic species that may affect the catalytic efficiency of the PETase when applied directly in *in situ* conditions (Zhu *et al*., 2021; Bergeson *et al*., 2024). To develop a potent enzyme-based technology, the effective enzymes capable of degrading PET plastics should be analyzed for their degradation efficiency in *in situ* conditions.

Phthalate diester plasticizers, including the high-priority pollutant bis(2-ethylhexyl) phthalate (DEHP), are extensively used in plastic products and accompany the increased plastic waste in landfills (Klotz *et al*., 2024). Previous studies have reported EstS1 esterase, a thermostable and pH-tolerant esterase, from *Sulfobacillus acidophilus* DSM10332 for its broad substrate specificity against a wide range of low molecular weight phthalate diesters plasticizers. The effective thermostability of this enzyme at a temperature above 70 °C provides it with an exceptional advantage (Zhang *et al*., 2014). Earlier, we discovered the degradation potential of this enzyme against the high molecular weight phthalate diester DEHP (Verma *et al*., 2024)

The present study elucidates the PET plastic-degrading ability of the EstS1 esterase, highlighting its potential to degrade both PET plastic and phthalate diester plasticizers. The biochemical analysis using HPLC and LC-MS was performed to reveal effective BHET degradation at 37 °C and formation of MHET, terephthalic acid, and ethylene glycol, disclosing the potential of EstS1 to mediate the conversion of BHET into MHET and then MHET to terephthalic acid. The crystal structure of wild-type EstS1 in complex with BHET revealed the electron densities of one molecule each of BHET, MHET, and ethylene glycol in the difference Fourier map, while the structure of Ser154Ala mutant of EstS1 esterase complexed with BHET showed intact two molecules of BHET tracing the active site tunnel of the enzyme, demonstrating the catalytically active form of the enzyme in the crystallization condition. Further, the molecular dynamics of EstS1-BHET complex over a period of 1 μs exposed the stable interaction of BHET at the catalytic site. Scanning Electron Microscopy (SEM) analysis of crystalline PET film showed significant degradation over the period of 15 days by EstS1 esterase. X-ray photoelectron spectroscopy (XPS) analysis revealed increase in -C-O-, -C-N-, - N-H-, and -N=O- bonds after degradation. Moreover, to mimic the environmental condition, the EstS1 PET degrading ability was evaluated in soil soaked with EstS1 esterase, and effective degradation was observed in ecological temperature conditions, proving the immense ability of EstS1 to withstand the environmental changes. Therefore, the present analysis determines EstS1 esterase as a double-edged sword capable of dealing with both plastic and plasticizer pollution.

## 2. Results

The exponentially growing pollution of plastics is posing harm to all life forms. The frequently used PET plastic accounts for a major portion of the total accumulated plastics on Earth in different environmental niches. Bioremediation has been identified as the most effective approach to deal with growing plastic levels (Joseph *et al*., 2024). Numerous PET-degrading enzymes have been characterized, and some engineered enzymes have been developed. Though much research has been conducted, very few effective enzymes for the degradation of crystalline PET plastic have been translated into technology for application in *in situ* conditions, either due to low efficiency against high crystallinity PET or temperature sensitivity, or inhibition from ionic species present in the environmental soil in *in situ* conditions (Samak *et al*., 2020). In the present study, EstS1 esterase, a thermostable and pH-tolerant phthalate diesters-degrading esterase, has been characterized for its potential to degrade PET plastic in controlled as well as in situ conditions, thus can be utilized for the development of technologies for bioremediation of both plastics and their plasticizers (Zhang *et al*., 2014; Verma *et al*., 2024).

### 2.1. BHET degradation kinetics of EstS1 esterase

The wild-type and Ser154Ala mutant of EstS1 esterase (34 kDa) cloned in the pET28a vector were purified up to 99% purity and concentrated to a concentration of 50 mg/mL (Figure S1). HPLC-based steady-state kinetics was performed using different concentrations of BHET to analyze the BHET degradation efficiency of the wild-type EstS1 esterase. The analysis of BHET in the presence and absence of EstS1 esterase was done for each considered concentration. The absorbance of BHET was monitored at 240 nm, and a significant peak of BHET was obtained at a retention time (t_R_) of 1.67 min (Figure 2). The reduction in the peak absorption in the presence of EstS1 esterase compared to the control was quantified using peak area, and thus, the degradation of BHET was estimated (Figure 2). The BHET degradation was observed to increase with the increase in concentration of BHET from 65% to 75%, indicating the first-order zone of the Michaelis-Menten kinetics, which encourages the most accurate estimation of k_cat_/K_m_. The k_cat_/K_m_ value estimated for BHET degradation was 13.63 M^-1^ sec^-1,^ and the V_max_ and K_m_ values were obtained to be 209.48 µM/min and 17.09 mM, respectively.

**Figure 2:**
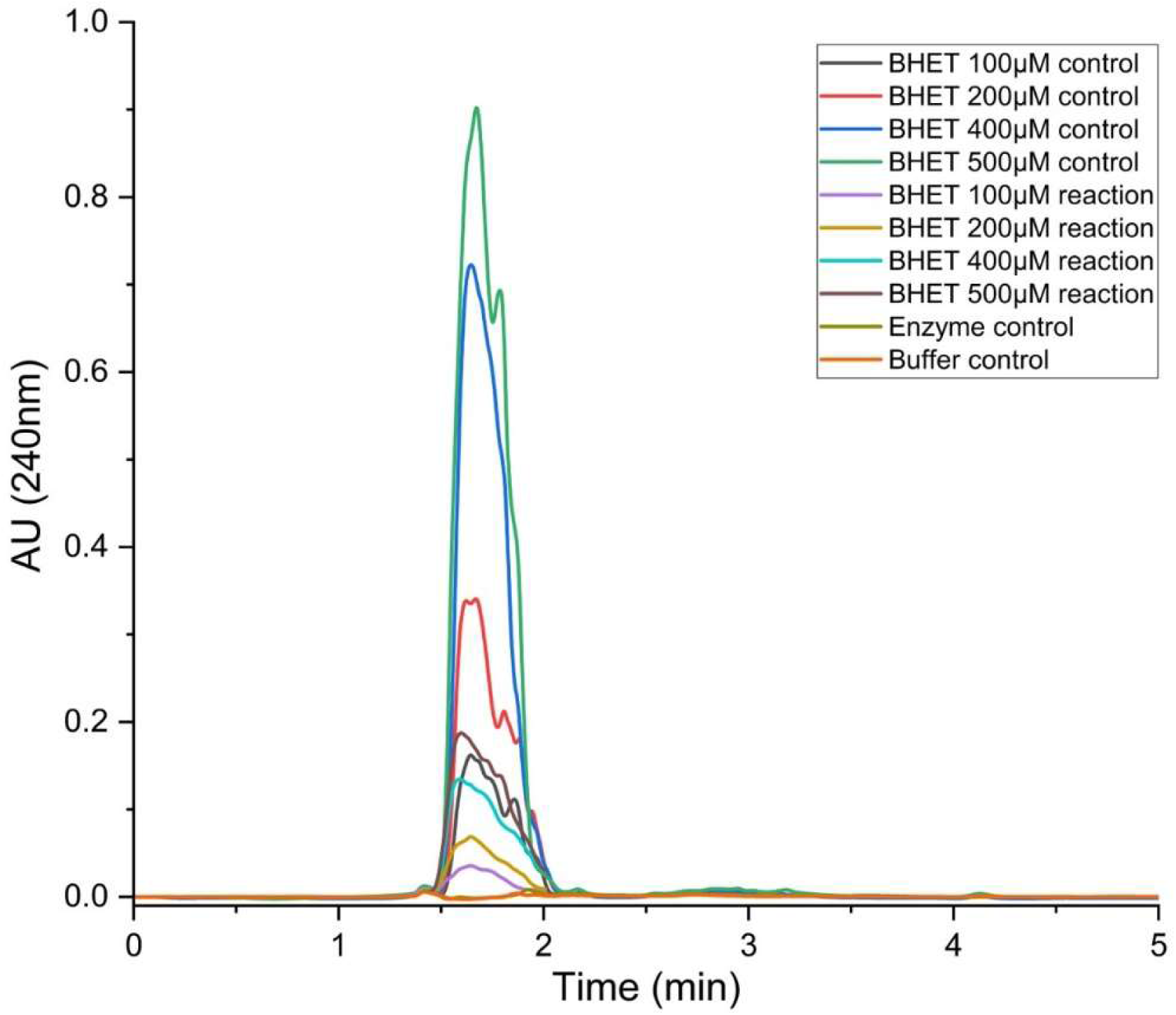
HPLC chromatogram showing bis(2-hydroxyethyl)terephthalate (BHET) degradation by EstS1 esterase. The absorbance at 240 nm showed a reduction in reaction (which corresponds to BHET concentration) as compared to the standard (control).

### 2.2. Identification of BHET degradation products

An LC-MS-based analysis was performed to further analyze the degradation of BHET by EstS1 esterase. The products of BHET degradation, i.e., MHET and ethylene glycol, along with TPA, were detected at retention times (t_R_) 8.12 min, 7.16 min, and 1.66 min, respectively, in negative ion mode (Figure 3 A, B, and C). Similar studies have reported the identification of the BHET products in negative ion modes, while BHET was identified in positive ion mode (Świderek *et al*., 2023) (Figure S2). The detection of TPA as a degradation product revealed the ability of EstS1 to catalyze the conversion of BHET to MHET and MHET to TPA, thus mediating cleavage of both the ester bond present in BHET (De Queiros Eugenio *et al*., 2021). The intensity versus mass/charge (m/z) ratio spectra of MHET, TPA, and ethylene glycol depict significant intensity peaks at the m/z values of 209.05, 165.02, and 62.96, which are close to the molecular masses of MHET (209.17 g/mol), TPA (164.11 g/mol), and ethylene glycol (62.07 g/mol) further confirming the presence of these compounds (Aristizábal-Lanza *et al*., 2022) (Figure 3 D, E, and F).

**Figure 3:**
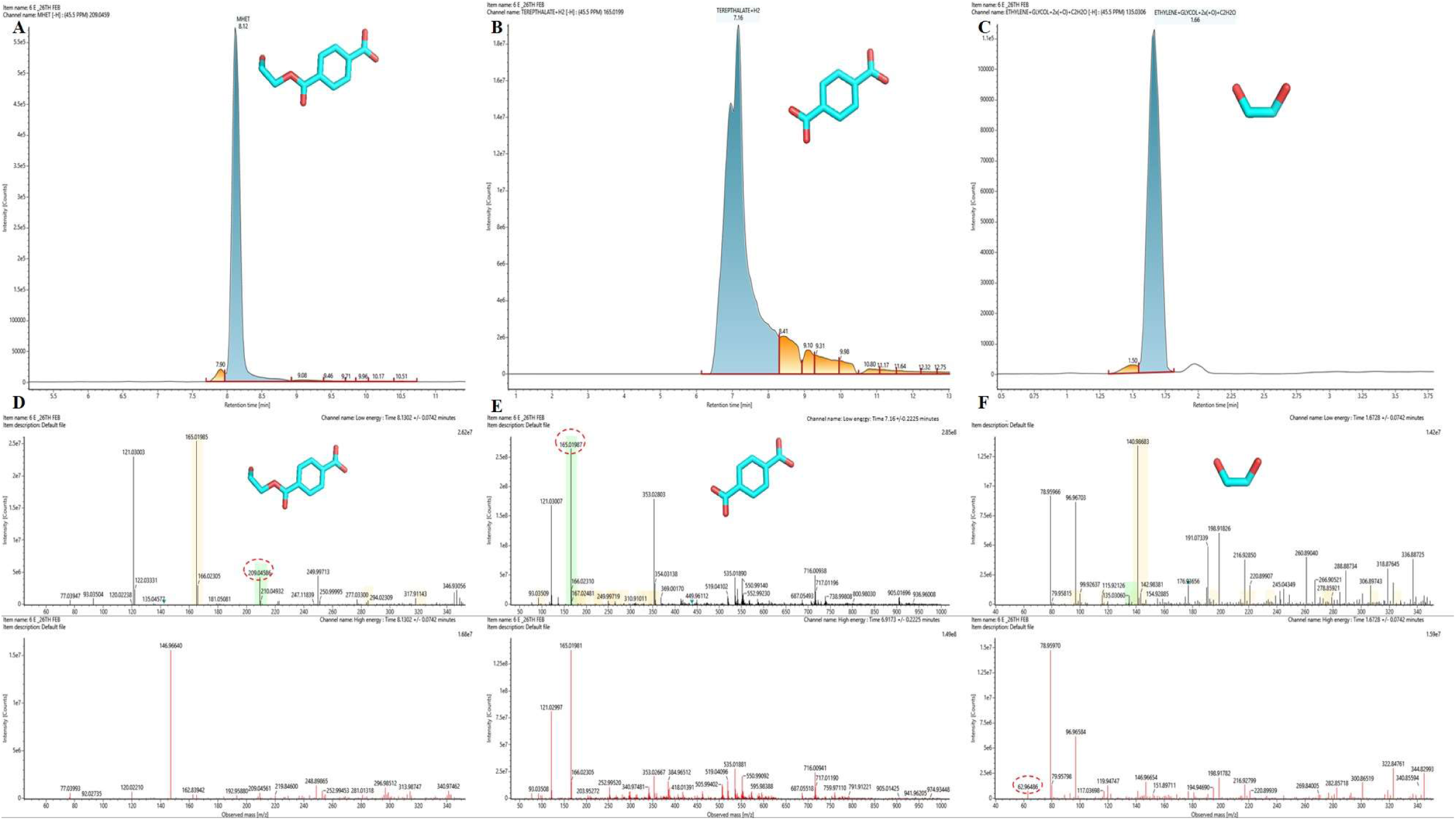
Liquid chromatography mass spectrometry (LC-MS) analysis of the bis(2-hydroxyethyl) terephthalate (BHET) degradation by EstS1 esterase. Intensity versus retention time chromatogram of BHET products A) mono(2-hydroxyethyl) terephthalate (MHET), B) terephthalate and C) ethylene glycol. Intensity versus observed mass (m/z) spectra at low and high energy of D) mono(2-hydroxyethyl) terephthalate (MHET), E) terephthalate, and F) ethylene glycol.

### 2.3. Complex crystal structures of wild-type EstS1 esterase with BHET

The co-crystal structure of wild-type EstS1 with BHET was solved in space group P63 at 2.2 Å with one monomer in the asymmetric unit. The refinement quality parameters of R*_work_* and R*_free_* were 19.2% and 24.95% (Table 1). The structure of EstS1 esterase comprised of 304 residues of which 12 residues (Glu8, Lys32, Arg70, Arg133, Lys145, Ser164, Arg166, Arg198, Ser231, Arg262, Glu269, Gln270) were observed in alternate conformation. The electron density map of residues Gln13 to Ser23 was missing due to the high flexibility of this loop. The EstS1 belongs to an α/β hydrolase family having a characteristic fold of 8 β-strands with intermittent 12 α-helices and a β-strand Pro60-Tyr66 running anti-parallel to the other 7 β-strands. The residues Tyr181 to Pro239 form the cap domain over the catalytic site, consisting of Ser154, His278, and Asp248 catalytic triad (Verma *et al*., 2024).

**Table 1:** Data collection and the refinement statistics of the Ser154Ala mutant EstS1-BHET complex and wild-type EstS1-BHET-MHET complex crystal structures.

| Parameter | EstS1-Ser154Ala_mutant-BHET (PDB ID: 9WNY) | Wild-type EstS1-BHET-Products (PDB ID: 9WNZ) |
| --- | --- | --- |
| Resolution range | 21.16 - 2.0 (2.072 - 2.0) | 21.21 - 2.2 (2.279 - 2.2) |
| Space group | P 63 | P 63 |
| Unit cell | 106.50 106.50 44.94 90 90 120 | 106.75 106.75 44.95 90 90 120 |
| Total reflections | 299692 (22317) | 121655 (9642) |
| Unique reflections | 19887 (1961) | 15058 (1484) |
| Multiplicity | 15.1 (11.4) | 8.1 (6.5) |
| Completeness (%) | 99.89 (100.00) | 99.84 (100.00) |
| Mean I/sigma(I) | 31.93 (6.97) | 31.14 (9.16) |
| Wilson B-factor | 11.1 | 10.8 |
| R-merge | 0.11 (0.38) | 0.1 (0.22) |
| R-meas | 0.11 (0.4) | 0.10 (0.24) |
| R-pim | 0.03 (0.12) | 0.033 (0.09) |
| CC1/2 | 1 (0.91) | 0.99 (0.92) |
| CC* | 1 (0.98) | 1 (0.98) |
| Reflections used in refinement | 19886 (1963) | 15057 (1486) |
| Reflections used for R-free | 1008 (123) | 770 (86) |
| R-work | 0.18 (0.19) | 0.19 (0.24) |
| R-free | 0.22 (0.22) | 0.25 (0.31) |
| CC(work) | 0.95 (0.88) | 0.94 (0.74) |
| CC(free) | 0.93 (0.82) | 0.87 (0.58) |
| Number of non-hydrogen atoms | 2453 | 2508 |
| macromolecules | 2322 | 2404 |
| ligands | 36 | 42 |
| solvent | 95 | 62 |
| Protein residues | 295 | 297 |
| RMS(bonds) | 0.01 | 0.011 |
| RMS(angles) | 1.93 | 2.12 |
| Ramachandran favored (%) | 97.25 | 96.25 |
| Ramachandran allowed (%) | 2.75 | 3.41 |
| Ramachandran outliers (%) | 0 | 0.34 |
| Rotamer outliers (%) | 3.28 | 7.2 |
| Clashscore | 8.14 | 13.04 |
| Average B-factor | 12.49 | 12.23 |
| macromolecules | 11.78 | 11.69 |
| ligands | 48.28 | 44.24 |
| solvent | 16.1 | 11.72 |

- Values in bracket represent last resolution shell values

The co-crystal structure of wild-type EstS1 with BHET revealed the catalytically active form of the enzyme in the crystal, as the difference Fourier map displayed the densities of one molecule each of BHET, MHET, and ethylene glycol contoured at 3σ (Figure 4 A and B). The overall structure of EstS1 consists of two surface cavities connecting together to form a tunnel that centres the catalytic site (Figure 4 C). The surface cavity 1, constructed by the cap domain heading toward the active site, has been identified in our previous work to mediate the entry of the substrate (Verma *et al*., 2024). In the present study, one molecule of MHET, the product of the reaction, was observed at the active site, forming hydrogen bonds with the catalytic Ser154, the oxyanion hole residues Ala155, Gly84, Gly83, and a water molecule, depicting the actual second tetrahedral intermediate of the canonical catalytic mechanism of esterase (Verma *et al*., 2024; Long & Cravatt, 2011) (Figure 4 B and C). The O4 atom of MHET formed a hydrogen bond with the OG atom of Ser154 at a distance of 2.5 Å, while the O5 atom of MHET formed 3 hydrogen bonds, with the backbone N of Ala155 at a distance of 3.3 Å, the oxygen atom of water at 2.7 Å, and the backbone N atom of Gly84 at 3.1 Å, respectively. The water molecule found in the catalytic site in turn, formed a hydrogen bond network with the catalytic serine and the oxyanion hole residues, thereby stabilizing the complex (Figure 4 B). The tail containing the second ester linkage of the formed product MHET was observed to be sandwiched between the two methionine residues Met186 and Met207, of which Met207 has been previously mentioned to stabilize substrate binding in the active site cavity to mediate the catalysis (Verma *et al*., 2024) (Figure 4 B). These two methionine residues contribute to the cap domain, which is likely to provide substrate specificity. Two water molecules were also present in the cavity, mediating the interaction of the O3 atom of the carbonyl carbon of MHET with the methionine residues (Figure 4 B). The BHET molecule in the catalytic tunnel was observed near cavity 2, due to the presence of hydrophilic residues Thr92, His93, and Thr282 in its close vicinity, the resulting orientation placed BHET at a distance from the catalytic triad and the oxyanion hole, thus preventing its degradation (Figure 4 B and C). Therefore, confirming cavity 1 as an effective site for entry of the substrate to mediate the catalysis.

**Figure 4:**
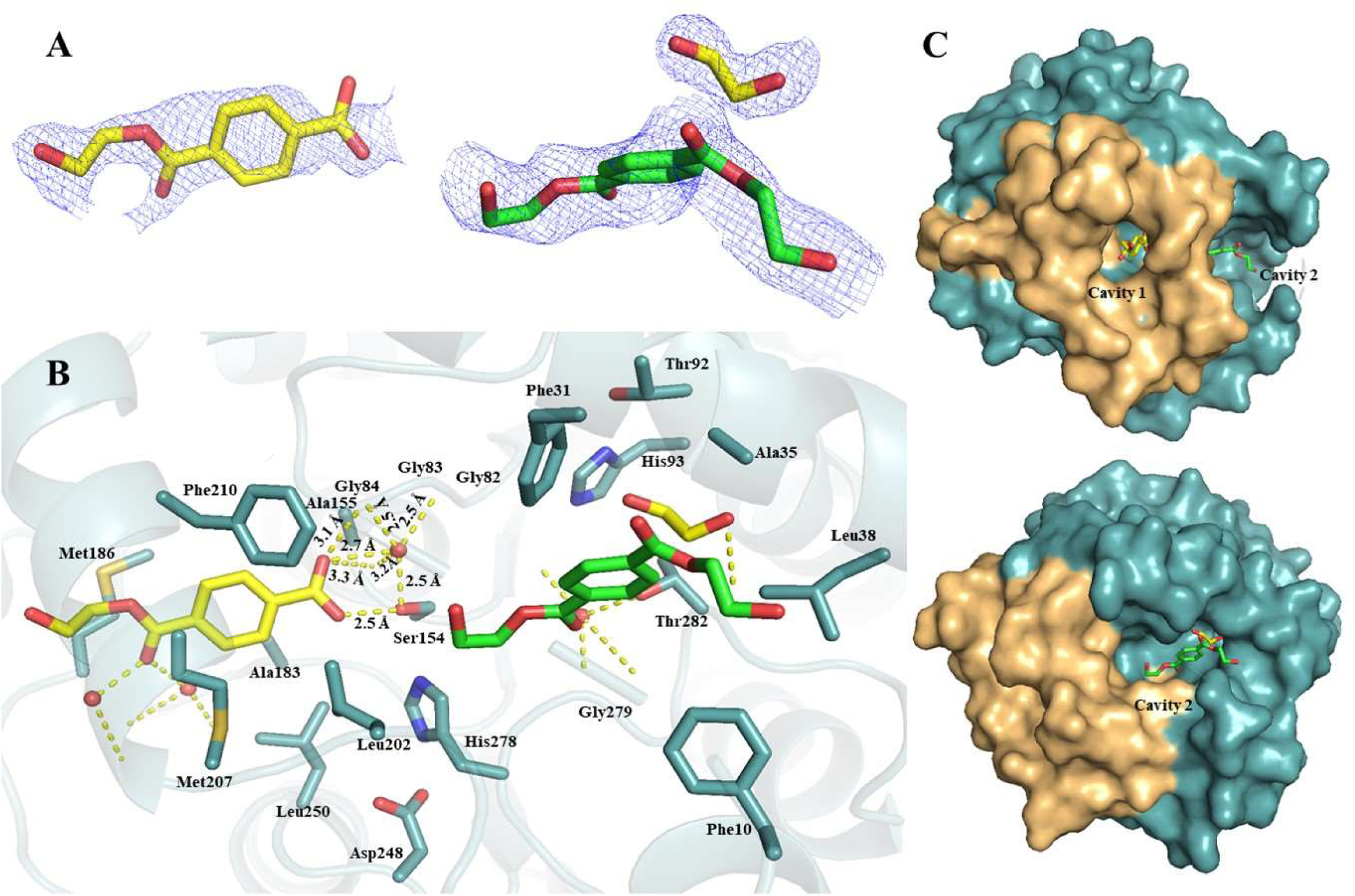
Crystal structure of wild-type EstS1 esterase with BHET and its degradation products. A) Electron density F_o_-F_c_ map contoured at 3σ of substrate BHET (green) and products, MHET (yellow), and ethylene glycol (yellow). B) Stick representation of EstS1 interactions with the substrate BHET (green) and products, MHET (yellow) and ethylene glycol (yellow) in the catalytic tunnel, showing interaction of MHET (Popout view) with the catalytic Ser154 and oxyanion hole residues. C) Surface representation of EstS1 esterase showing the relative positions of BHET and MHET inside the catalytic tunnel. (Yellow dotted lines displaying hydrogen bond interactions).

### 2.4. Complex crystal structures of Ser154Ala mutant EstS1 esterase with BHET

To further understand the interaction of EstS1 esterase with BHET an active site mutant of Ser154Ala EstS1 was purified and co-crystallized with the BHET. The co-crystal structure of Ser154Ala mutant EstS1 esterase with BHET was solved at 2 Å in P63 space group with R*_work_* and R*_free_*of 17.81% and 22.16% respectively (Table 1). The residues Arg6, Lys32, Thr57, Ile129, Lys168, Ser222, and Ser231 were found to be present in alternate conformation, and the electron density of the loop comprising Gln13 to Ser23 residues was missing. This complex structure revealed the electron densities of two BHET molecules in the difference Fourier map contoured at 3σ (Figure 5 A). One molecule of BHET was observed at the active site and seemed to enter from cavity 1, while the other molecule was in front of the first one and towards cavity 2 (Figure 5 B). The orientation of the BHET molecule bound at the active was similar to that of the degraded product MHET discussed above and was favourable for mediating catalysis but due the presence of alanine (Ala154) in place of catalytic serine (Ser154) the carbonyl carbon of the ester bond did not interact with the Ala154 while the terminal hydroxyl group O1 atom of the BHET formed hydrogen bonds with backbone nitrogen atoms of Gly83, Gly84 and Ala155 at a distance of 2.8, 2.7 and 2.9 Å respectively. Moreover, the hydrophilic residues and network of water in the catalytic tunnel towards cavity 2 resulted in a curved orientation of the second BHET molecule, resulting in its inaccessibility to the catalytic site (Figure 5 B and C).

**Figure 5:**
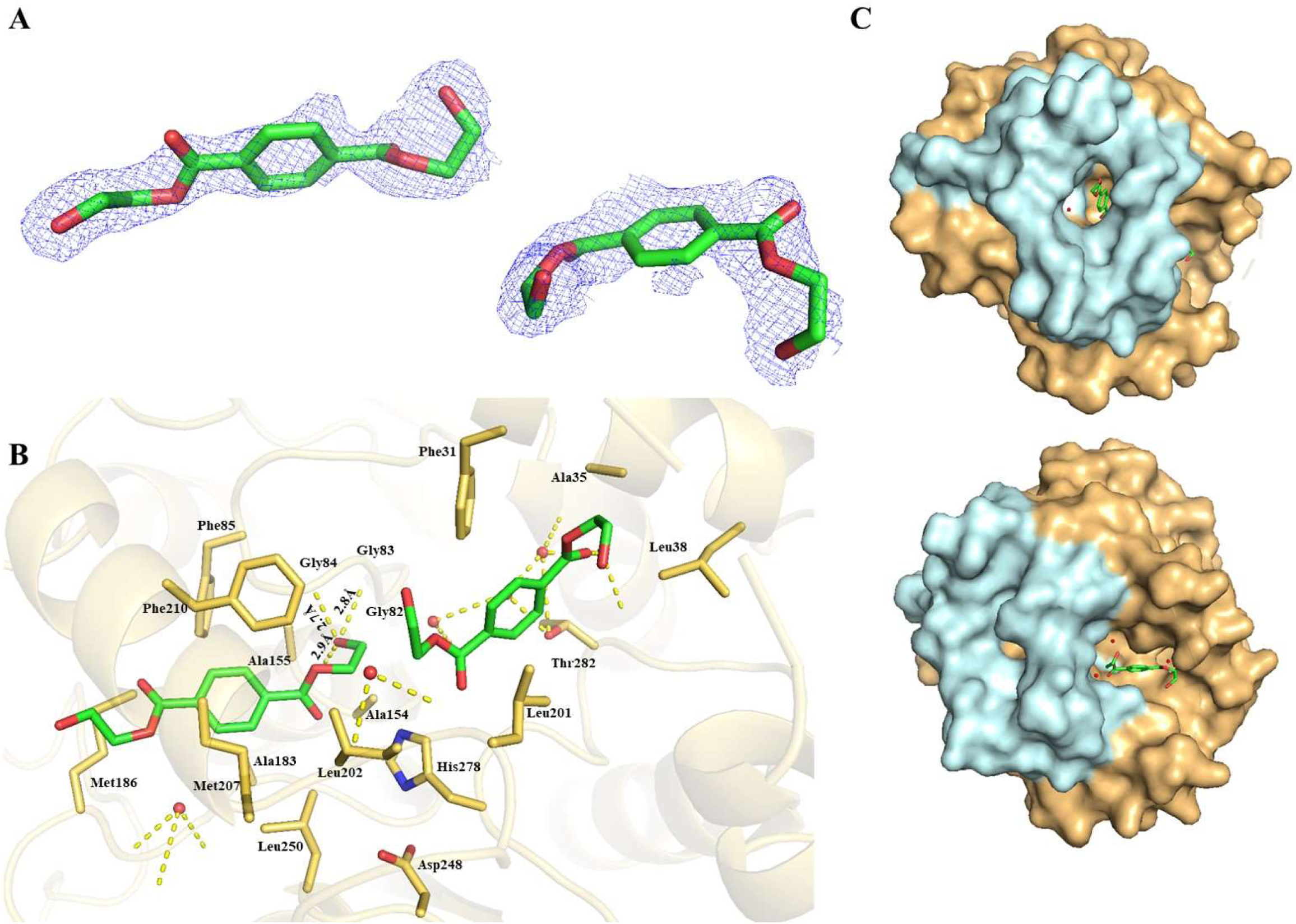
Crystal structure of Ser154Ala mutant of EstS1 esterase with BHET. A) Electron density F_o_-F_c_ map contoured at 3σ of 2 molecules of substrate BHET (green). B) Stick representation of EstS1 esterase with 2 molecules of BHET (green) in the catalytic tunnel. C) Surface representation of EstS1 esterase showing the relative positions of 2 molecules of BHET inside the catalytic tunnel. (Yellow dotted lines displaying hydrogen bond interactions).

Therefore, structural analysis of EstS1-BHET complexes revealed that though BHET seems to enter from both the cavity 1 (hydrophobic) and 2 (hydrophilic) due to the presence of hydrophobic ring and carbon chain, and the hydrophilic terminal hydroxyl groups respectively but the hydrophobic nature of the cavity 1 mediated binding of BHET in the proper orientation of the hydrophobic ring and the carbon chains tail to promote catalysis.

### 2.5. Dynamics of EstS1-BHET interactions

Molecular dynamics (MD) simulation of apo-EstS1 and EstS1-BHET complex to study the interaction dynamics of EstS1 esterase with BHET was performed over a period of 1 μs. The analysis of the MD simulation trajectory revealed high stability of the EstS1 esterase when present in complex form as compared to the apo-protein. The RMSD values ranged within a narrow range of 0.20 to 0.30 nm in complex form compared to 0.15 to 0.35 nm for the apo-form of the protein, indicating high stability of the complex (Schreiner *et al*., 2012) (Figure 6 A). The least variation in Rg values within 1.81 to 1.85 nm disclosed high compactness of the protein during the simulation trajectory of 1 μs (Filipe & Loura, 2022) (Figure 6 B). The RMSF versus residue plots showed a reduction in fluctuation in residues ranging from Tyr181 to Pro239 in complex form compared to the apo-protein. These residues contribute to the cap domain, which mediates the entry of the substrate. The reduction in fluctuations in the cap domain in complex form discloses that binding of the substrate at the catalytic site imparts stability to the cap domain during the simulation (Benson & Daggett, 2012) (Figure 6 C). The solvent accessible surface area analysis during the trajectory time showed variation in the range of 127 to 140 nm^2^ in complex form, while the variation in apo-form ranged from 123 to 144 nm^2^, in turn showing reduction in solvent accessible surface area and hence increase in compactness and stability of the protein in complex form (Durham *et al*., 2009) (Figure 6 D). The variation in secondary structures showed an increase in the alpha-helicity and a significant decrease in the coiled structures in the complex form compared to the apo-form of the enzyme, indicating high stability of the complex form (Coskuner-Weber & Caglayan, 2021) (Figure 6 E and F). The steepness of the free energy landscape of EstS1-BHET complex revealed attainment of global minima or a properly folded low-energy state during the trajectory, as compared to local minima or partially folded states observed in the apo-protein (Maisuradze *et al*., 2010) (Figure 7 A and B). The consistent intermolecular hydrogen bonding between EstS1 and BHET showed effective interaction during the trajectory at the active site of the enzyme (Bondar & White, 2011) (Figure 7 C). The clustering analysis of the 1 μs trajectory disclosed a large number of structures in the top 10 clusters (Figure 7 D). The superimposition of the representative structures of the top 10 clusters revealed that the BHET was consistently bound at the active site, interacting with the catalytic triad and the oxyanion hole residues (Phillips *et al*., 2011) (Figure 7 E). Thus, the present analysis provides compelling evidence for the highly stable and consistent interaction of BHET at the active site of EstS1 esterase.

**Figure 6:**
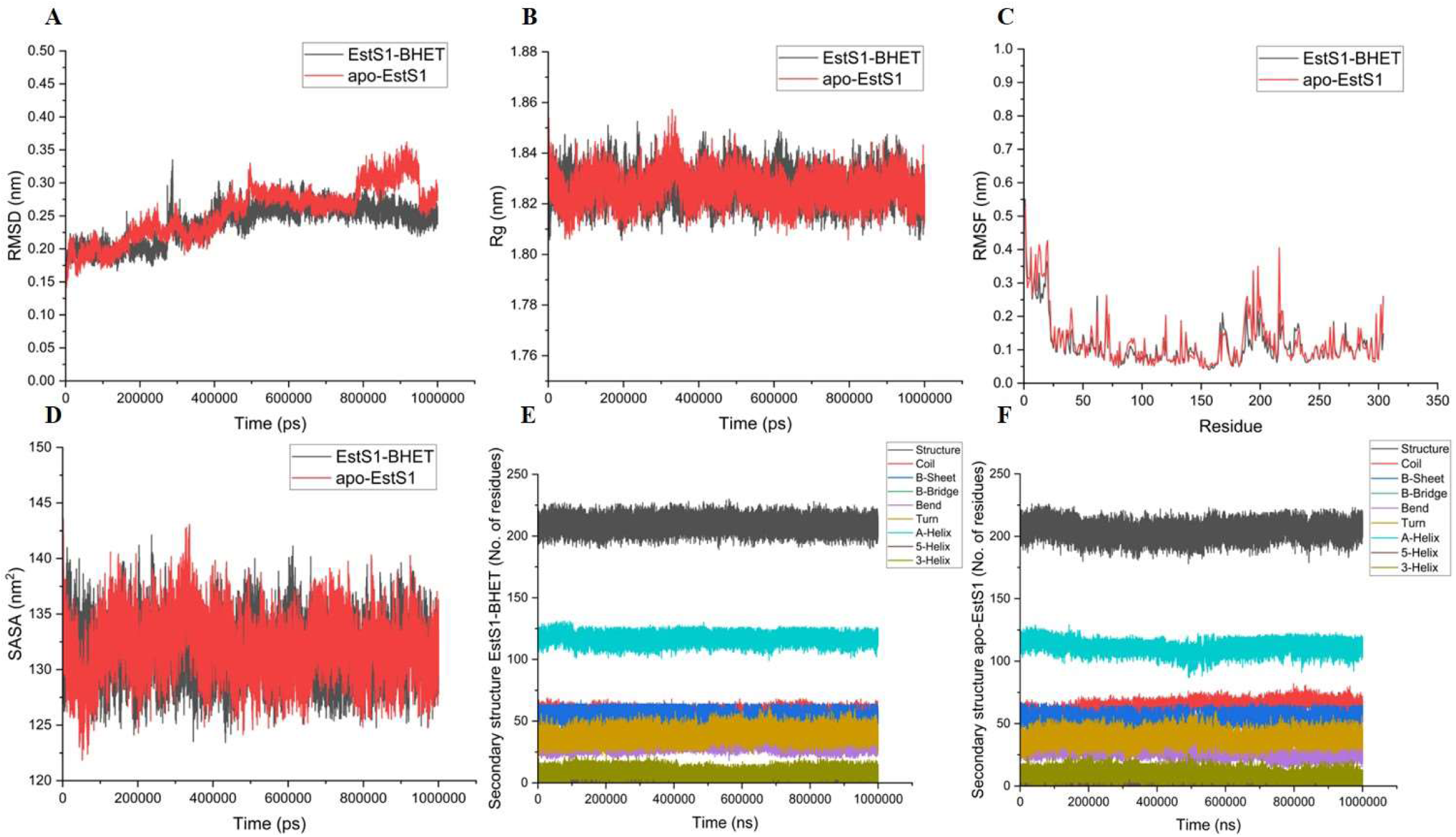
Molecular Dynamics Simulation analysis of the EstS1-BHET complex and apo-EstS1 esterase over a trajectory time of 1 μs. A) Root Mean Square Deviation (RMSD) of protein backbone versus time plot. B) Radius of gyration (Rg) of protein backbone versus time plot. C) Root Mean Square Fluctuation (RMSF) versus residue plot. D) Solvent Accessible Surface Area (SASA) versus time plot. E) Line plot of secondary structure (SS) composition of EstS1-BHET complex versus trajectory time. F) Line plot of secondary structure (SS) composition of apo-EstS1 versus trajectory time.

**Figure 7:**
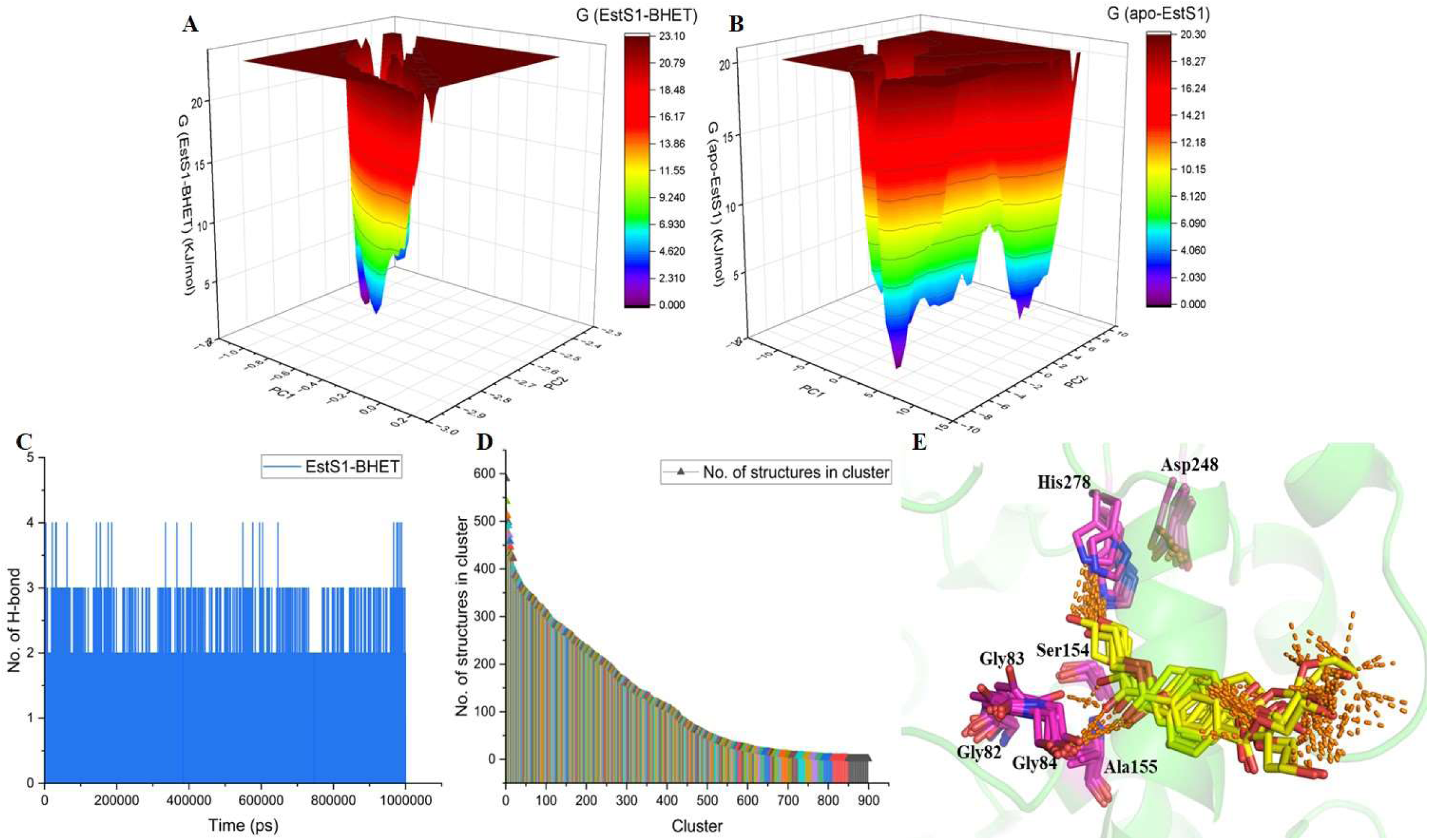
(Continued) Molecular Dynamics Simulation analysis of the EstS1-BHET complex and apo-EstS1 esterase over a trajectory time of 1 μs. A) Free energy landscape of EstS1 esterase in the presence of BHET. B) Free energy landscape of apo-EstS1 esterase. C) Number of hydrogen bonds formed versus trajectory time plot of EstS1-BHET complex. D) Bar plot of the number of structures versus clusters generated in clustering analysis of EstS1-BHET complex. E) Superimposition of representative conformations of the top 10 clusters of EstS1-BHET complex.

### 2.6. Microscopic analysis of PET film for degradation in controlled conditions at 25 °C and in environmental conditions in soil

The biochemical, crystal structure, and molecular dynamics simulation-based analysis disclosed the efficiency of EstS1 esterase in degrading BHET, a monomer of PET film. To evaluate the PET film degrading efficiency of the EstS1 esterase, the crystalline PET film was soaked with the EstS1 esterase for a duration of 15 days in controlled temperature conditions at 25 °C. The SEM analysis of the PET film after a duration of 15 days displayed erosion and cracks on PET film, revealing efficient EstS1-mediated degradation of crystalline PET film visible at a 1000X to 10000X magnification (Figure 8). Further to analyze the efficiency of PET degradation in natural environmental conditions, the PET film was incubated in soil augmented with EstS1 esterase and soil without the enzyme as a control sample. The samples were placed in a natural environment in glass bottles to encounter regular variations in temperature and pressure for 15 days. The SEM analysis of the samples after 15 days showed indents and erosion of the surface of the film, concluding efficient degradation of the PET film at 1000X to 25000X magnification in the presence of diverse ionic concentrations of soil (Figure 9). This analysis provides evidence for the significant applicability of the EstS1 esterase in *in situ* conditions to ameliorate the increasing concentration of microplastics and plastics in the environment.

**Figure 8:**
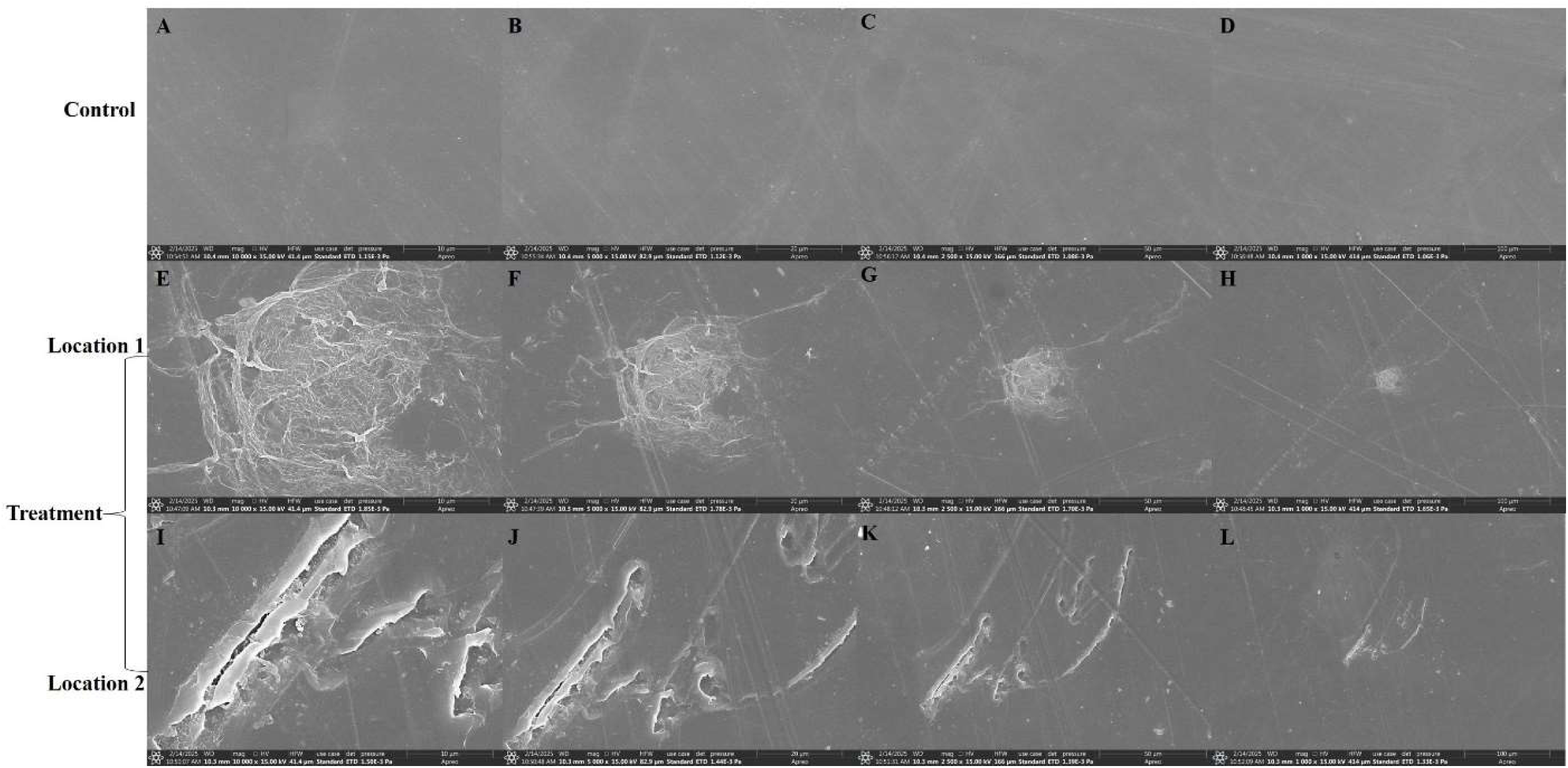
Scanning electron microscopy analysis of crystalline PET film degradation by EstS1 esterase at 25 °C over 15 days. A-D) SEM images of the control sample (PET film soaked in buffer) at 10000X, 5000X, 2500X, and 1000X magnification. E-H) SEM images of the treatment sample (focused at one location in PET film soaked in EstS1 esterase) at 10000X, 5000X, 2500X, and 1000X magnification, showing erosion of the PET film. I-L) SEM images of the treatment sample (focused at another location in PET film soaked in EstS1 esterase) at 10000X, 5000X, 2500X, and 1000X magnification, showing indents and erosion of the PET film.

**Figure 9:**
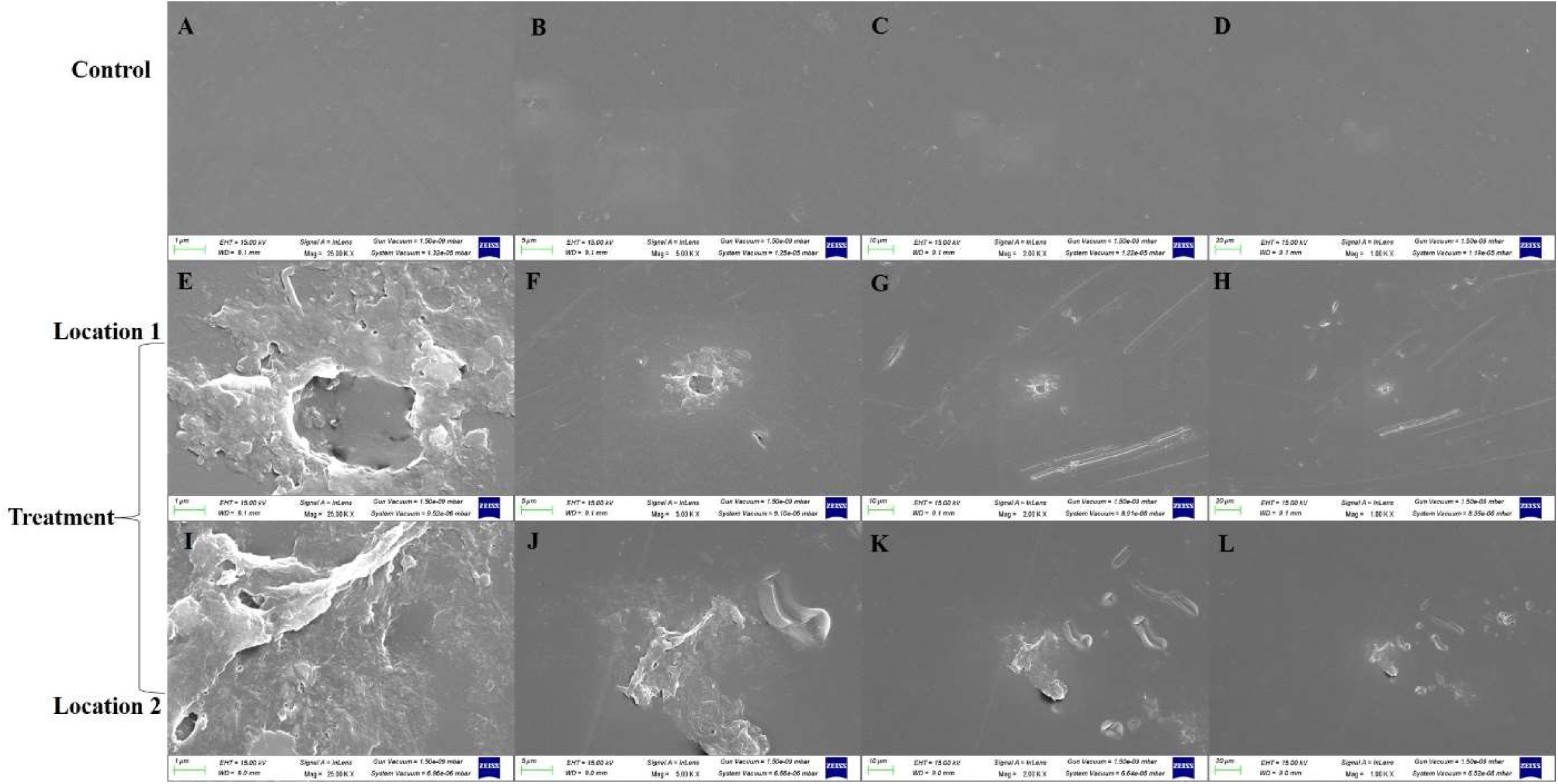
Scanning electron microscopy analysis of crystalline PET film degradation by EstS1 esterase in the soil microenvironment exposed to environmental variation of temperature. A-D) SEM images of the control sample (PET film soaked in soil) at 10000X, 5000X, 2500X, and 1000X magnification. E-H) SEM images of the treatment sample (focused at one location in the PET film in soil containing EstS1 esterase) at 10000X, 5000X, 2500X, and 1000X magnification, showing indents and erosion of the PET film. I-L) SEM images of the treatment sample (focused at other location in the PET film in EstS1 esterase) at 10000X, 5000X, 2500X, and 1000X magnification, showing indents and erosion of the PET film.

### 2.7. Elemental and surface chemistry analysis of PET film for degradation by EstS1

XPS analysis of the PET film sample subjected to EstS1 enzyme treatment and the control sample was performed to evaluate the changes in elemental composition and properties of chemical bonding due to degradation by EstS1. The counts per second versus binding energy survey spectra of surface (initial) and at a depth of 50 nm (final), displaying the atomic composition, reveal the presence of nitrogen element in the treated sample compared to the absence of nitrogen in the control sample (Figure 10 A B). The concentration of nitrogen in treatment sample was higher at the surface than at a depth of 50 nm, therefore confirming the presence of nitrogen-containing degraded products similar to the past study (Huang et al., 2024) (Figure 10 A B). The curve fitted C1s intensity versus binding energy spectra of both control (C) and treated (T) samples were analysed and significant peaks at 284.562 eV and 287.876 eV were obtained in control sample which corresponds to -C-C-, and -C=O- bond while peaks at 284.562 eV, 285.776 eV and 287.876 eV were observed in treatment sample corresponding to -C-C-, -C-O-, -C-N-, -C=N- and -C=O- bonds disclosing the formation of -C- O- and -C-N- bonds after degradation (Figure 10 C).

**Figure 10:**
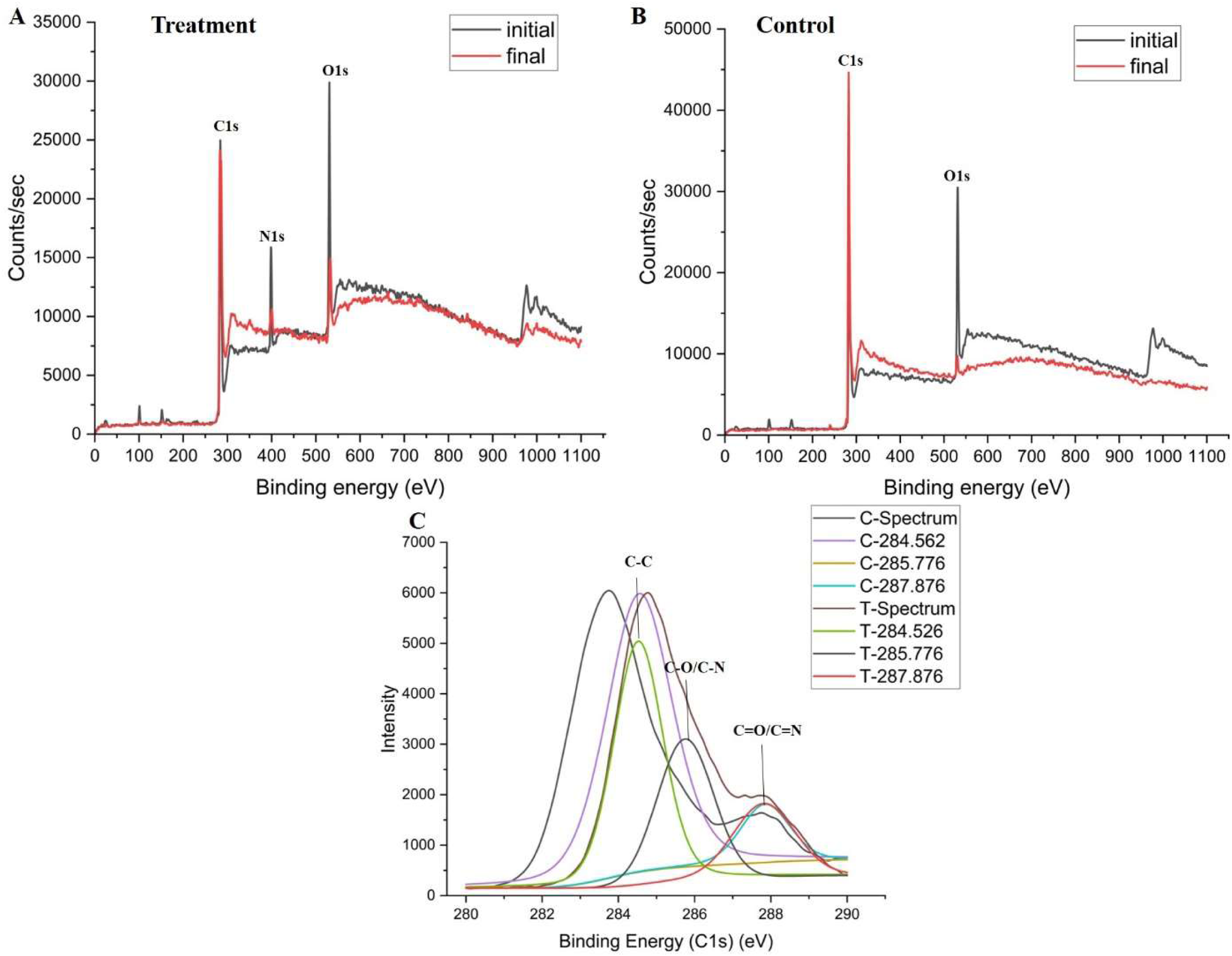
XPS survey spectra of upper surface and final surface at a depth of 50 nm of A) Treatment sample and B) Control sample. C) Comparison of peak fits of high resolution C1s spectra of upper surface of treatment (T) and control (C) samples.

The depth analysis of the treated and control PET film for variation in chemical bonding properties at every 3 nm up to 50 nm depth was done. The intensity versus binding energy spectra of C1s, N1s, and O1s showed significantly high presence of C-O-, -C-N-, -N-H- (400 eV), -N-O- (404 and 408 eV), and -C-O- bonds indicated by peaks at different binding energies in treated samples with higher prevalence at the upper surface compared to the inner surface (Figure 11 A B C). In control samples, the peaks were consistent within a short binding energy range, showing less diversity in chemical bonding patterns over 50 nm depth (Figure 11 D E F). The analysis of variation in atomic concentration with respect to depth of film showed a high amount of nitrogen and oxygen at the surface in the treatment sample at the upper surface compared to the control sample. The carbon content at the surface of the treated sample was less in comparison to the control sample surface, indicating erosion of surface (Figure 11 G).

**Figure 11:**
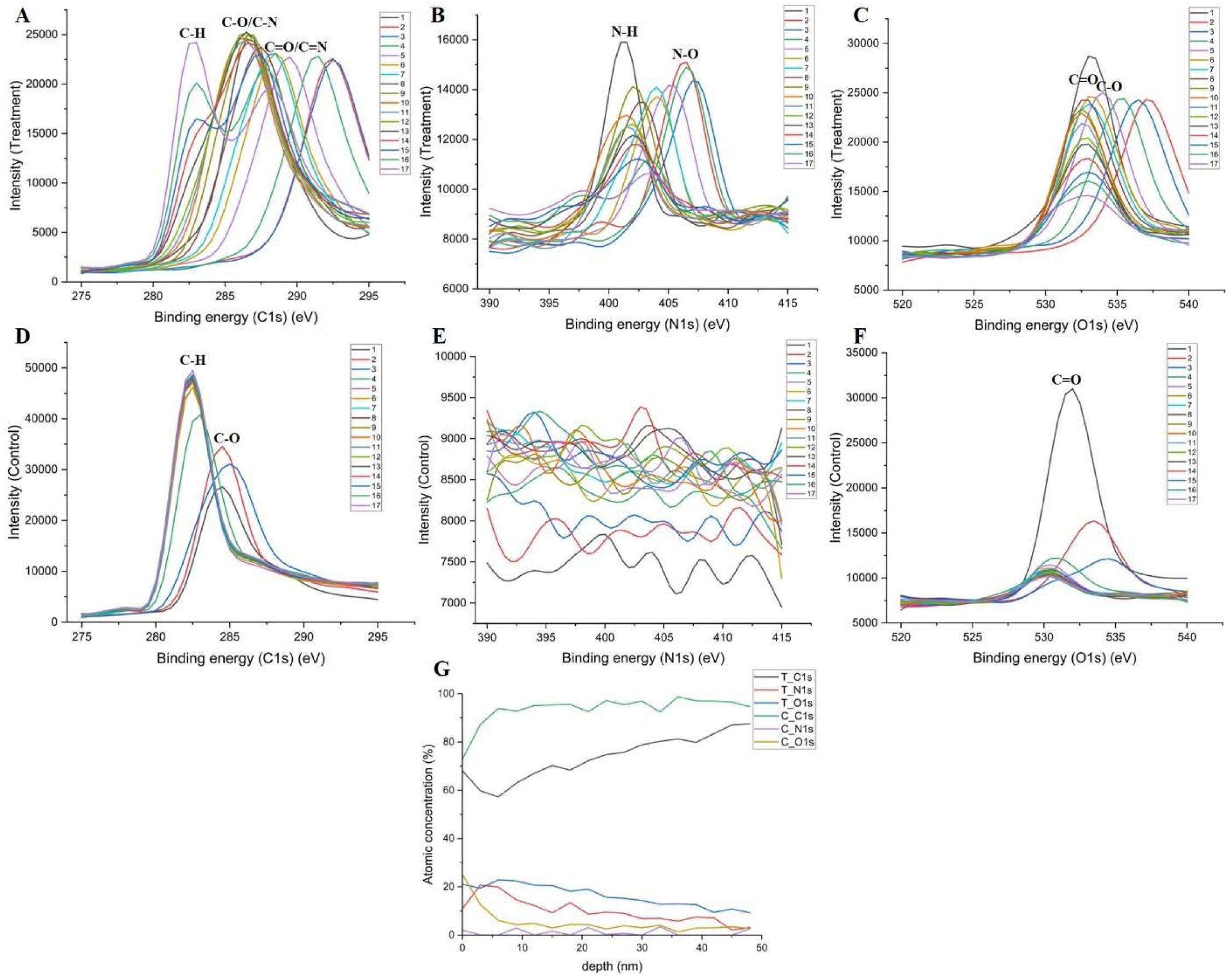
XPS high resolution spectra of C1s N1s and O1s of A-C) treatment and D-F) control samples over a depth of 50 nm (1-17 indicate surface at every 3 nm over the range of 0 nm to 50 nm) G) XPS atomic concentration profiles of treatment (T) and control (C) samples over a depth of 50 nm.

## 3. Discussion

Enzymatic degradation of PET plastic is being intensely investigated to deal with the major fraction of accumulated PET plastic in the environment, which is responsible for increasing plastic pollution. Diverse PETases, BHETases, cutinases, lipases, and esterases have been studied for their efficiency in degrading different PET varieties, including amorphous PET, nanoPET, semi-crystalline PET, and crystalline PET. Several studies have engineered these enzymes to enhance their thermostability and efficiency to a greater extent (Bell *et al*., 2022; Cui *et al*., 2024; Shi *et al*., 2023; Pfaff *et al*., 2022).

EstS1 esterase from *Sulfobacillus acidophilus* DSM10332 strain is a thermostable, pH-tolerant phthalate diester-degrading enzyme capable of degrading a broad range of phthalate diesters (Zhang *et al*., 2014; Verma *et al*., 2024). The present study elucidates the BHET and PET degrading ability of the EstS1 esterase utilizing biochemical, structural, *in silico*, and SEM analysis-based approach. Around 75% degradation of BHET by EstS1 esterase was observed over a period of 1 h, resulting in the formation of MHET and TPA, concluding the ability of EstS1 esterase to convert BHET into MHET and then MHET into TPA. Previous study on *Streptomyces* strains characterized the BHET-degrading potential of *Streptomyces* LipA lipase variants and identified mild TPA formation compared to MHET, with low activity above 40 °C due to the lack of enzyme stability at high temperatures (Verschoor *et al*., 2024). EstS1 esterase, a highly thermostable esterase, annotated as triacylglycerol lipase, produced a significant concentration of TPA as compared to the MHET. *Ht*DLH, a thermostable hydrolase of *Hydrogenobacter thermophilus* reported for PET, BHET, and MHET degradation, shows comparable BHET degradation to EstS1 esterase (Almeida *et al*., 2024). Several other studies reported engineered PETase enzymes having high catalytic activity against BHET, while the catalytic efficiencies of wild-type enzymes were comparable to that of EstS1 esterase analyzed in the present study (Bell *et al*., 2022; Pfaff *et al*., 2022; Numoto *et al*., 2020; Kitadokoro *et al*., 2012; Sulaiman *et al*., 2011).

To illustrate the structural aspects of BHET interaction at the active site of EstS1 esterase, the co-crystal structure of wild-type EstS1 esterase with BHET was determined and revealed not only the presence of the active form of EstS1 esterase in the crystal form, but also the density of MHET and ethylene glycol along with BHET. The bound MHET (the product of BHET degradation) at the active site depicted the exact second tetrahedral intermediate conformation of the canonical serine hydrolase mechanism. The cap domain, which mediates the substrate specificity, varies in length in different esterase and PETase enzymes (Distaso *et al*., 2023). The cap domain of EstS1 of 59 residues forms cavity 1, which mediates the entry of the substrate. The two methionine (Met186 and Met207) residues of the EstS1 esterase, lying in the cap domain, provide a proper orientation to the substrate to interact with the catalytic triad, thus mediating catalysis. Similar contributions of the cap domain residues in mediating effective substrate binding have been reported in previous studies on MHETase (Knott *et al*., 2020). Joo *et al*., 2018, in their structural analysis of *Is*PETase, reported Met161 and Ile208 as the lead residues providing a hydrophobic surface for substrate binding, similar to the hydrophobic residues Met186 and Met207 present in cavity 1 of the EstS1 esterase (Joo *et al*., 2018). The presence of one intact BHET molecule (undegraded) near cavity 2, even in the active form of the enzyme, in turn determines the essentiality of the cap domain (forming cavity 1) in contributing to the catalysis by providing correct orientation to the substrate at the active site. Comparative analysis of known PETase enzymes with EstS1 disclosed the least sequence and structural similarity, revealing a novel PET-degrading enzyme scaffold in EstS1 esterase. The large cap domain of EstS1, compared to the other PET-degrading enzymes, provides a massive scope for further enzyme engineering studies to improve its efficiency (Luan *et al*., 2016).

The complex structure of the Ser154Ala active site mutant of EstS1 in complex with BHET exhibited the presence of two BHET molecules bound in the catalytic tunnel of EstS1, one after the other, demonstrating the significant size of the tunnel to accommodate the PET polymer. Further, *in silico* structural analysis displayed high stability of the EstS1 esterase in the presence of BHET, directing towards highly favorable interactions of BHET with the active site residues of the enzyme. The free energy landscape of the EstS1-BHET complex showed attainment of the stable low-energy state of the enzyme (Benkovic *et al*., 2008).

Moreover, the erosion and indentation of crystalline PET film surface in the presence of the enzyme in both controlled conditions and soil system with varying temperature conditions confirmed effective crystalline PET degradation potential, high stability, and ionic tolerance of EstS1 esterase. Though several studies have analyzed the PET film degradation using amorphous PET film, nano PET, and PET microplastics, very few studies have been conducted for the degradation of crystalline PET in conditions mimicking the original environment (Wilkes *et al*., 2024; Ibrahim *et al*., 2024; J. Kim *et al*., 2023). The XPS-based surface and depth profiling analysis discovered a higher concentration of nitrogen in the treatment sample compared to the extremely low concentration in the control samples. Also, the formation of -C- O-, -C-N-, -C=O-, -C=N-, -N-H-, and -N=O- bonds was observed to increase in the treatment sample while the prevalence of these bonds was less in the control samples, thus proving the degradation of the PET film (Doren et al., 1994). Further, the atomic concentration versus depth plot showed a lower proportion of carbon on the surface of the enzyme-treated sample compared to the control sample in turn indicating erosion of the surface. Similar increase in nitrogen concentration in enzyme treated PET film have been observed in past studies indicating effective degradation and production of nitrogen conjugated products containing -C- N-, -C=N-, -N-H- and -N=O- bonds along with increased formation of -C-O-, and -C=O- bonds was reported (Huang et al., 2024; Han et al., 2024)

Therefore, the present study elucidates the PET-degrading potential of a high-potential thermostable EstS1 esterase. Its efficiency in degrading crystalline PET plastic and its monomer BHET have been evaluated, along with its in-depth structural analysis, displaying the mechanism of degradation. Hence, it can be further exploited for enzyme engineering studies to develop into a high-efficiency PET as well as phthalate diester plasticizer-degrading enzyme, providing a complete solution to the increasing plastic and plasticizer pollution.

## 4. Conclusion

Global plastic pollution has drawn widespread concerns due to its adverse impacts on all life forms across the globe. PET-based plastics are leading contributors to the increasing plastic pollution. Bioremediation of plastics using potential enzymes is a safe, eco-friendly, and effective approach. In this piece of work, we have identified the BHET and PET degrading potential of a EstS1 phthalate diesters-degrading pH-tolerant and thermostable esterase from *Sulfobacillus acidophilus* DSM10332 strain. The kinetic and LC-MS-based analysis disclosed up to 75% degradation of BHET, a PET monomer, into MHET and terephthalic acid. Complex crystal structures of wild-type and mutant forms of EstS1 esterase with BHET have demonstrated the mechanistic insights of the enzyme, representing the cap domain residues as effective contributors responsible for the proper orientation of the substrate at the active site. Further, the cavity 1 formed by the cap domain was identified as the exact entry site for the substrate to mediate the reaction. *In silico* analysis of the EstS1-BHET complex exhibited that the enzyme attains a highly stable, low-energy folded state in the presence of substrate. Eventually, the variations in elemental composition and chemical bonding via XPS and corroded surface and indents on the surface of crystalline PET film via SEM analysis were observed in incubated PET film with EstS1 esterase, which provided strong evidence for the PET degradation potential of EstS1 esterase in constant and varying temperature conditions, mimicking the environmental variations. Henceforth, further enzyme engineering of EstS1 esterase can provide a highly potent solution to the exponentially growing plastics as well as plasticizer pollution to attain the goals of a sustainable environment.

## 5. Materials and Methods

### 5.1. Cloning, Expression, and Purification of EstS1 and Its Active Site Mutant

The cloning, expression, and purification of EstS1 esterase and its mutant were performed according to our previously mentioned procedure (Verma *et al*., 2024). The pET28a recombinant plasmid of EstS1 was transformed into *E. coli* BL21 (DE3) cells (Microbial Type Culture Collection (MTCC) Chandigarh, India). The 1 L Luria-Bertani liquid broth was inoculated at 1 % v/v inoculum and kept at 37 °C until the optical density of the culture at 600 nm reached 0.6. The expression was checked by adding isopropyl β-D-thiogalactopyranoside (IPTG) as inducer at a concentration of 0.5 mM. The induced culture was shifted to 16 °C for incubation for 16 h. The cell mass was obtained and resuspended in 10 mL lysis or purification buffer (100 mM phosphate buffer, 10 % v/v glycerol, 2 mM DTT, pH 7.4) at 4 °C. The cells were lysed at 20 psi using a constant cell disruptor, and the lysate was centrifuged for 45 min at 12000 rpm at 4 °C. The supernatant was collected and transferred to a pre-equilibrated Ni-NTA affinity column with purification buffer and incubated for 10 min (Mahto *et al*., 2022). The gradient of imidazole from 10 to 500 mM was applied to the incubated Ni-NTA column, and the fractions were collected. SDS-PAGE was performed to analyze the fractions, and the fractions containing pure protein were pooled and concentrated using a 30 kDa Millipore Amplicon filter and were later dialyzed with 100 mM HEPES buffer at pH 7.4 (Verma *et al*., 2024). The concentration of the protein was evaluated using bovine serum albumin (BSA) standard and Pierce Micro BCA protein assay kit (Thermo Scientific).

A site-directed mutagenesis approach was used to prepare the active site mutant Ser154Ala of EstS1 esterase (Bachman, 2013). The recombinant plasmids of pET28a-EstS1 were amplified with forward and reverse primers as mentioned below. The product of amplification was subjected to DpnI enzyme digestion, and then transformed into *E. coli* DH5α cells. The transformed cells were cultured, and the mutated plasmids were isolated using the Plasmid Isolation Kit (Qiagen QIAprep Spin Miniprep Kit Cat no. 27104) and confirmed by DNA sequencing (Verma *et al*., 2024). The transformation of the confirmed plasmids was done in *E. coli* BL21 (DE3) cells, and the mutant protein was purified using the above-mentioned protocol.

*Forward Primer*: GTGGCCGGGGACGCGGCCGGCGGCAATTTG

*Reverse Primer*: GCCGCCGGCCGCGTCCCCGGCCACCACAATC

### 5.2. HPLC analysis of BHET degradation kinetics

The BHET degradation was quantified using a Waters high-performance liquid chromatography (HPLC). The reactions of 1 mL each containing different concentrations (100, 200, 400, and 500 μM) of BHET were incubated with 15 μM of enzyme for 1 h at 37 °C. The termination of reactions was done by the addition of 100% methanol in a 1:1 ratio, and the reactions were then filtered through nylon syringe filters (0.2 μm) (Li *et al*., 2023). The BHET degradation was analysed on a Waters C18 XBridge column (150 x 4.6 mm; particle size 5 µ) by isocratic elution (0.7 mL/min), using a mobile phase consisting of methanol–acidified water (0.1% formic acid) in the ratio of 80:20 (Lyoo *et al*., 2000; Boros *et al*., 2025). The chromatograms were monitored at 240 nm. The buffer and enzyme control samples were also analyzed, along with the BHET standard solutions (100, 200, 400, and 500 μM) without enzyme to plot the standard curve. All control and enzyme-treated samples were analyzed in triplicate, and average values were utilized for the calculation of V_max_, K_m,_ and k_cat_/K_m_ (Counotte & Prins, 1979).

### 5.3. LC-MS analysis for product identification

Liquid chromatography-mass spectrometry (LC-MS) was performed to analyze the degradation of BHET by EstS1 esterase. The 1 mL enzyme reactions containing 5 mM BHET and 15 μM EstS1 esterase were incubated at 37°C for 1 h in triplicate. The reaction was terminated by adding 1mL 100% methanol. The reactions were then filtered through 0.2 μm nylon filters. Chromatographic separation using reverse phase chromatography was performed using a C18 reverse phase column (Waters ACQUITY UPLC BEH column) of dimension 1.7 μm, 2.1 mm × 100 mm. The mobile phase consists of 0.1 % v/v formic acid in methanol and water at a flow rate of 0.3 mL/min. The elution gradient was used over 20 min, ranging from 100% water to 100% methanol (Swamy *et al*., 2024).

Mass spectrometry analysis was done using an Xevo G2-XS QTof Quadrupole Time-of-Flight Mass Spectrometry system (Waters) in both positive and negative ionization modes. The conditions applied for mass spectrum acquisition consist of cone and capillary voltages of 50 V and 3 kV, respectively. The source and desolvation temperatures were 450 °C and 120 °C, and the gas flow rates of the cone and desolvation were 50 L/h and 900 L/h, respectively. Electron spray ionization with a scan range of 100-1000 m/z and 1 s scan time was used. The parameter of collision energy for low energy was 6 eV, and for high energy, a ramp from 20 to 30 eV was set with a time of acquisition of 8 min. Processing of LC-MS data was done using Waters UNIFI software (Waters), and the target by mass approach was utilized. Other parameters, including the maximum number of targets, fragment match tolerance, target match tolerance, and ion ratio tolerance, were specified. Further threshold values from candidates per sample, relative intensity, and peaks per channel were added. The expected ratio values of m/z for BHET, MHET, terephthalic acid, and ethylene glycol were identified by MestReNova software (https://mestrelab.com/) for use in preparing the library.

### 5.4. Co-crystallization of EstS1 wild type and EstS1 Ser154Ala with BHET

The co-crystallization of both wild-type and Ser154Ala active site mutant of EstS1 at an enzyme concentration of 10 mg/mL was performed with BHET (10 mM) using our previously mentioned crystallization condition of 100 mM HEPES pH 7.0, 0.5% v/v Jeffamine ED-2001, and 1.1 M sodium malonate. The sitting drop method for crystallization was used at a 1:1 protein to reservoir ratio and a temperature of 20 °C. The rod-shaped crystals obtained were cryoprotected in mother liquor and flash-frozen in a liquid nitrogen stream at 100 K (Verma *et al*., 2024).

Data collection was performed at the home source X-ray diffractometer integrated with an X-ray generator (Rigaku Micromax-007 HF), having a high-intensity rotating microfocus Cu anode and a Hypix Rigaku 6000C detector available at the Macromolecular Crystallography Unit, IIC, IIT Roorkee. The complex crystal structures were solved using the apo-EstS1 structure (PDB: 9J1E) as search model using the molecular replacement method MOLREP of CCP4i2 suite and Phenix 1.20.1 (Verma *et al*., 2024; Vagin & Teplyakov, 2009; Potterton *et al*., 2018; Adams *et al*., 2010). Iterative rounds of building the model, followed by refinement, were performed utilizing Coot v0.9.8.95, REFMAC, and phenix.refine (Emsley & Cowtan, 2004; Murshudov *et al*., 2011; Afonine *et al*., 2012). The co-crystal structure of the wild-type EstS1-BHET complex revealed the presence of BHET as well as MHET and ethylene glycol in the catalytic tunnel, while the co-crystal Ser154Ala active site mutant of EstS1-BHET resulted in electron densities of two molecules of BHET in the catalytic tunnel. Omit maps were generated for the bound ligands using Phenix. The interaction and surface diagrams were generated using PyMol (https://www.pymol.org/) and Chimera (https://www.cgl.ucsf.edu/chimera/) (Rigsby & Parker, 2016; Pettersen *et al*., 2004).

### 5.5. Molecular dynamics (MD) simulation analysis

The co-crystal structure of Ser154Ala mutant EstS1-BHET was used to study the dynamics of interaction of BHET within the active site tunnel using the molecular dynamics simulation approach. The alanine residue at the active site was mutated to a serine residue using Coot software to analyze the actual mechanism of interaction of wild-type EstS1 with BHET. The apo and the BHET complex forms of EstS1 were subjected to 1 μs MD simulations each using the GROMACS 2020 software (Pronk *et al*., 2013). The CGenFF web server was used to generate the topology of BHET, and the CHARMM36 force field was used for the topology of the protein (Vanommeslaeghe *et al*., 2009; Huang & MacKerell, 2013). The dodecahedron box was used for solvation of the apo and complex structures of the EstS1 esterase, and the systems were neutralized using sodium ions. The energy minimization of the systems was performed using a maximum force of 10 kJ/mol. Then the minimized systems were equilibrated using Number of particles, Volume, Temperature (NVT) and Number of particles, Pressure, Temperature (NPT) ensembles for 100 ps each. Eventually, the MD simulation run of 1 μs each was performed for both the apo and BHET complex systems.

The analysis of the MD simulation trajectories was performed by evaluating variations in Root Mean Square Deviation (RMSD), Radius of gyration (Rg), Root Mean Square Fluctuation (RMSF), Solvent Accessible Surface Area (SASA), Secondary Structure (SS), and number of hydrogen bonds during the trajectory (Hollingsworth & Dror, 2018). The Principal Component Analysis (PCA) was performed for all the trajectory frames, and the free energy landscapes were generated for both the apo-protein and complex form of the protein (David & Jacobs, 2013). The clustering analysis to study the effective interactions of BHET with the catalytic tunnel of EstS1 was done considering an RMSD cutoff of 0.2, and the representative frames of the top 10 clusters were superimposed to study changes in the structure in complex form and the interaction mechanism of EstS1 with BHET (Abramyan *et al*., 2016). All plots of the MD simulation analysis were prepared using Origin 2024 software (https://www.originlab.com/), and images of representative clusters showing 3D interactions of BHET with EstS1 were generated using PyMol software (https://www.pymol.org/) (Rigsby & Parker, 2016).

### 5.6. PET film degradation analysis using Scanning Electron Microscopy (SEM) imaging

The small pieces of crystalline PET films (thickness 0.1 mm, Goodfellow) were soaked in the presence of EstS1 esterase (15 μM) for a duration of 15 days at 25 °C. Control samples without protein using 100 mM phosphate buffer at pH 7.4 were also prepared. After 15 days, the samples were collected and washed with 70% ethanol and distilled water to remove traces of enzyme and buffer. The morphological analysis of films was done for both enzyme-treated and control samples using a scanning electron microscope (ThermoFisher, Apreo S Low Vac, 2011) at 15 kV accelerating voltage. Film samples were gold-coated at 18 mA for 140 s. Digital images were generated using Apreo SEM imaging software (Yan *et al*., 2020; Ducoli *et al*., 2023).

### 5.7. Analysis of PET film degradation in natural conditions

The ionic composition of soil often poses an inhibitory effect on the enzymes present in the soil. To analyze the effect of soil ionic composition on EstS1 PET degradation activity, the PET film was cut into small pieces of equal weight and incubated in samples containing soil+buffer (soil control), and soil+buffer+EstS1 (soil treatment). The soil used for the experiment was collected from the nearest garden at IIC, IIT Roorkee, and the unwanted particles were removed. The clean soil was subjected to autoclaving at 15 psi for 20 min to remove the possibility of any other microbe enhancing the degradation. 6 g of autoclaved soil was used in each soil sample (treatment and control).15 μM of enzyme concentration was used in the treatment samples of soil, and an equivalent volume of buffer was used in the control sample. All control and treatment samples were kept outside in native environmental conditions in covered small glass bottles to mimic the *in situ* conditions of temperature (ranging from 40 °C in the afternoon to 28 °C at night) and pressure for 15 days (Liu *et al*., 2023). The degradation effect of EstS1 esterase in treatment versus control samples was analyzed using a scanning electron microscope (EVO 18, Carl Zeiss, 2012, and ZEISS SmartSEM software) at 15 kV as mentioned above. The PET film samples were washed with 70% ethanol before SEM analysis, followed by distilled water to remove any traces of soil and enzymes. All experiments were performed in triplicate. This experiment provides scope for the direct *in situ* application of the EstS1 esterase in soil for PET plastic degradation (Yan *et al*., 2020; Ducoli *et al*., 2023).

### 5.8. PET film degradation analysis by X-ray photoelectron spectroscopy (XPS) analysis

To further analyze the change in surface chemistry due to the degradation of the PET film by EstS1 esterase, the surface analysis of the PET film was performed by X-ray photoelectron spectroscopy. The PET film (thickness 0.1 mm, Goodfellow) samples incubated with EstS1 esterase (15 μM) for 15 days, along with control samples incubated with an equivalent volume of 100 mM phosphate buffer in natural environmental conditions in small glass bottles, were subjected to XPS analysis. The PET samples were washed with 70% ethanol and twice with distilled water to remove any traces of enzyme and buffer from the samples. The change in carbon, oxygen, and nitrogen elements’ composition, chemical state, and bonding was studied up to a depth of 50 nm by using X-ray photoelectron spectroscopy (PHI 5000 Versa Probe III; Ulvac-Phi, Inc). Ultrahigh vacuum of 2 * 10^-7^ Pa with Al Kα monochromatic source of energy 1.48 keV was used. The beam diameter of 100 µm, stage tilt angle of 45°, charge neutralizer with 15 V and 20 µA, and non-conductive tape as substrate were used for the analysis. Survey scan spectra were collected with an energy of 280 eV and 0.05 eV energy step. The high-resolution spectra of C1s, O1s, and N1s were recorded with 55 eV pass energy and 0.05 eV energy step. The C1s standard at 284.80 eV was considered as a reference. To analyze the chemical bonds, the carbon C1s and oxygen O1s region peaks were fitted. The Multipak XPS software for deconvolution and data analysis was used (Doren et al., 1994).

## Supporting information

concentrated to a concentration of 50 mg/mL (Figure S1).

9WNY

9WNZ

## Statements and Declarations

## ACKNOWLEDGEMENT

We thank the X-Ray Crystallography facility provided by MCU, IIC, IIT Roorkee, Ashok Soota Molecular Medicine Facility at Department of Biosciences and Bioengineering, IIT Roorkee for biochemical experimental work, the computational facility was provided by DBT funded project “Translational and Structural Bioinformatics – BIC at Department of Biotechnology, IIT, Roorkee (BT/PR40141/BTIS/137/16/2021) to P.K.

## DATA AVAILABILITY

The X-ray crystal structures of EstS1 esterase active site mutant S154A in complex with Bis(2-hydroxyethyl) terephthalate (BHET) and EstS1 esterase in complex with mono(2-hydroxyethyl) terephthalate (MHET) and bis(2-hydroxyethyl) terephthalate (BHET) have been deposited to the RCSB Protein Data Bank under PDB accession codes of 9WNY and 9WNZ respectively.

## AUTHOR CREDIT STATEMENT

Conceptualization (P.K., S.V.), Data curation (S.V., D.A., M.A., A.K.P., S.P.) Formal analysis (S.V., D.A., M.A., A.S., A.K.P.) Funding acquisition (P.K.) Investigation (S.V., D.A., M.A., AKP) Methodology (S.V., A.K.P., D.S., P.K.) Project administration (P.K.) Resources (P.K.) Software (S.V., A.K.P., P.K.) Supervision (P.K.) Validation (D.S., P.K.) Visualization Writing – original draft (S.V., P.K.) Writing – review & editing (D.S., J.S., P.K.)

## FUNDING AND FACILITIES

This work was funded by DBT-funded projects entitled “Structural insight into the molecular mechanism of enzymes involved in biodegradation of microplastics” (Grant No. BT/PR45880/BCE/8/1634/2022) to PK.

## ETHICS APPROVAL

Not applicable

## DECLARATION OF GENERATIVE AI

AI-based tool has not been used for this manuscript

