## Supplementary material for "Unveiling the PET plastics degradation potential of the thermostable EstS1 esterase through integrated biochemical, structural, and morphological analyses": concentrated to a concentration of 50 mg/mL (Figure S1).

***Running title:*** *PET plastics degradation by EstS1 esterase*

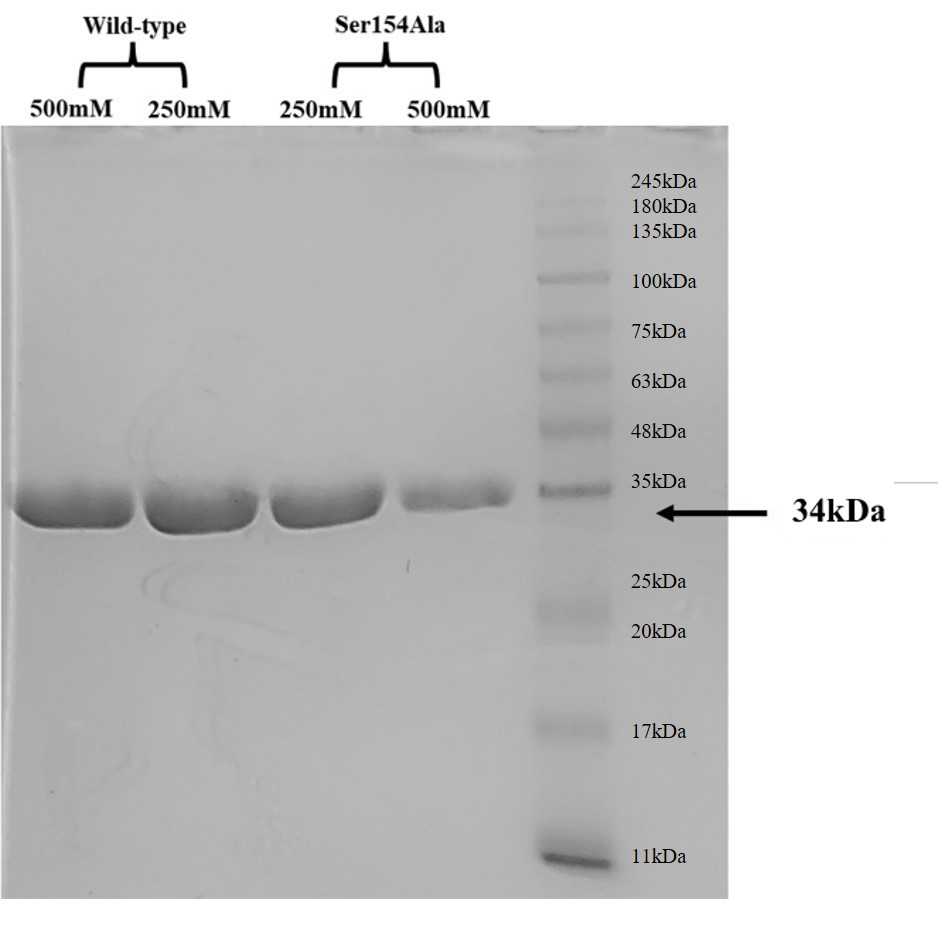

**Figure S1:** SDS PAGE gel image showing clear bands of wild-type and Ser154Ala mutant of EstS1 esterase at 34kDa

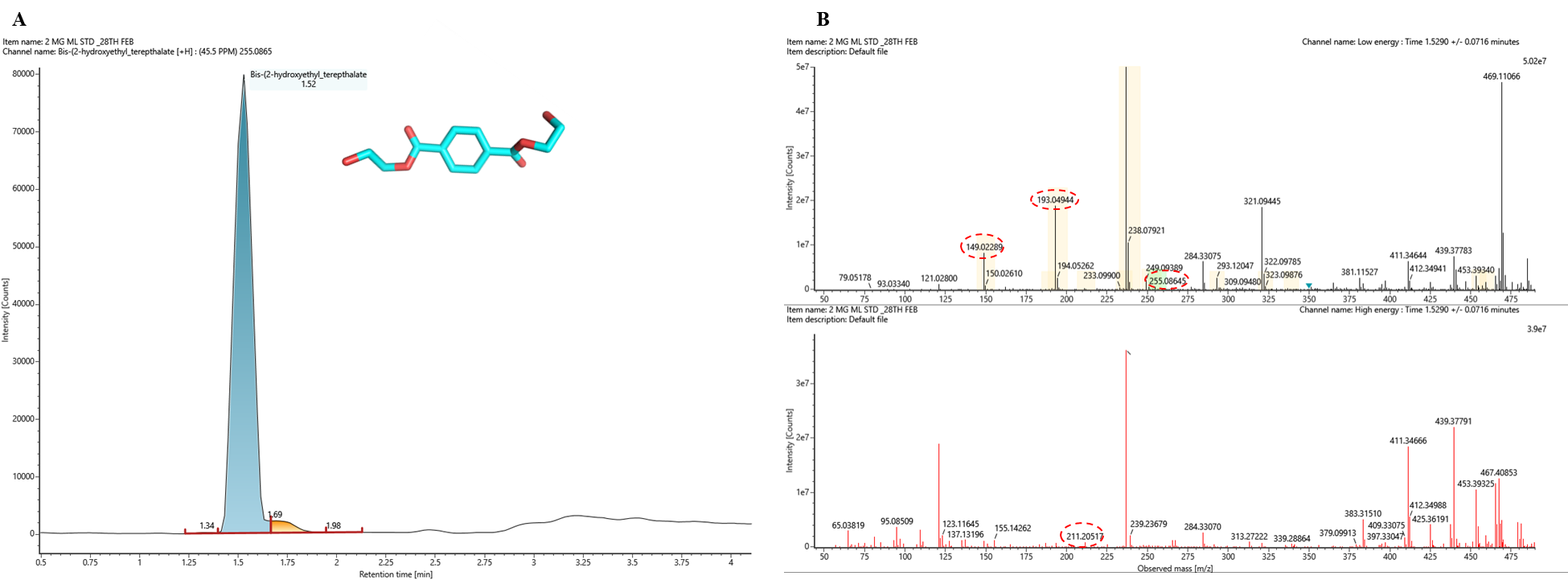

**Figure S2**: Liquid chromatography mass spectrometry (LC-MS) standard chromatogram and mass spectra of the bis(2-hydroxyethyl) terephthalate (BHET) in positive ion mode. A) Intensity versus retention time chromatogram of BHET. B) Intensity versus observed mass (m/z) spectra at low and high energy of BHET.
